# Animal chromosome counts reveal similar range of chromosome numbers but with less polyploidy in animals compared to flowering plants

**DOI:** 10.1101/2020.10.10.334722

**Authors:** Cristian Román-Palacios, Cesar A. Medina, Shing H. Zhan, Michael S. Barker

## Abstract

Understanding the mechanisms that underlie chromosome evolution may provide insights into the processes underpinning the origin, persistence, and evolutionary tempo of lineages. Here we present the first database of chromosome counts for animals (the Animal Chromosome Count database, ACC) summarizing chromosome numbers for ∼18,000 species. We found remarkable similarity in the distribution of chromosome counts between animals and flowering plants. At larger timescales, selection towards a specific range might explain the similar distribution of chromosome counts between these two groups. Nevertheless, changes in chromosome number are still potential drivers of divergence among species at shorter timescales. We also found that while animals and plants exhibit similar frequencies of speciation-related changes in chromosome number, plant speciation is more often related to changes in ploidy. Based on the ACC, our analyses suggest that changes in chromosome number alone could help explain patterns of diversity within animal clades.

## Introduction

The number of chromosomes in a nuclear genome is a fundamental aspect of eukaryotic biology. Differences in chromosome number and arrangement are also one of the longest studied topics in comparative genomics (e.g. Stebbins 1950; White 1973, 1978; Grant 1981; King 1995). Despite this long history, how and why chromosome numbers evolve remains poorly understood. One obstacle to improving comparative analyses and hypothesis testing is a lack of easily available chromosome counts from across the tree of life. Although many chromosome counts are available in the published literature (Peruzzi and Bedini 2014), these have only been made easily available for specific clades including plants (Rice et al 2015), coleoptera (Blackmon and Demuth 2015), polyneoptera (Sylvester and Blackmon 2020), amphibians (Perkins et al. 2019), mammals (Martinez et al. 2017; Blackmon et al. 2019), and fish (Arai 2011; Martinez et al.2015) along with other groups scattered across the Tree of Life (The Tree of Sex Consortium 2014). Analyses of these publicly available data have driven new understandings of chromosome and genome evolution (e.g. Zhan et al. 2014; Barker et al. 2016a; Salman-Minkov et al. 2016; Zenil-Ferguson et al. 2019). Increasing access to other publicly available chromosome counts from the literature will be important for understanding how and why chromosome numbers vary across the Tree of Life.

Here, we present the first database of chromosome counts from across all animal clades summarizing haploid and diploid counts for ∼18,000 animal species across 21 phyla. We used the current release of the ACC to examine fundamental questions about the distribution of chromosome counts and evolution of haploid numbers in animals. These included evaluating whether within-clade heterogeneity in chromosome number scales with clade age, testing if haploid chromosome counts are phylogenetically conserved, inferring the frequency of polyploidy, and estimating how often speciation is associated with changes in chromosome numbers or coupled with polyploidization. We also compared the patterns of chromosomal evolution in animals to those from angiosperms and ferns. We expect our newly compiled public animal chromosome counts database to fuel new comparative analyses that further our understanding of the forces driving chromosome evolution across the Tree of Life.

## Methods

We compiled animal chromosome counts from primary sources, books, and published datasets (e.g. Makino 1951; Benazzi and Benazzi 1976, along with other volumes of *Animal Cytogenetics*; O’Brien et al. 2006; Gokhman 2009; Arai 2011; Graphodatsky et al. 2012; Olmo et al. 2012; Ashman 2014; Blackmon et al. 2019; Perkins et al. 2019; Sylvester and Blackmon 2020). Sources for each chromosome count in the ACC are included in the database. We used the GBIF taxonomy backbone, accessed using the rgbif R package (Chamberlain et al. 2020), to curate the ACC.

We used phylogenetic and descriptive methods to examine the evolution of chromosome numbers across the animal Tree of Life. First, we used phylogenetic comparative methods to examine the evolutionary patterns of chromosome number evolution across the animal Tree of Life. We used phylogenetic regression models to test whether older phyla have accumulated more differences in chromosome numbers among species relative to younger phyla. Similarly, we tested whether diversity in chromosome numbers is related to species diversity across phyla. Clade ages, species richness, and phylum-level topologies follow Wiens (2015). Average differences in median chromosome counts were estimated as the mean pairwise difference in chromosome number among all species within each phylum. We fit phylogenetic regressions using the caper R package (Orme et al. 2013), with the Lambda parameter estimated from the data. Next, we tested for phylogenetic signal in chromosome numbers across species within orders sampled in the ACC. Time-calibrated species-level phylogenies within orders were retrieved from the TimeTree of Life database (Hedges et al. 2015). We estimated the phylogenetic signal in chromosome counts for animal orders with chromosome counts in the ACC and phylogenetic information from at least ten species. Phylogenetic signal in chromosome counts was estimated based on Pagel’s Lambda (Pagel 1999) using the phylosig function implemented in the phytools R package (Revell 2012). Finally, we examined whether the evolution of chromosome numbers within orders are best described by a neutral model of evolution (BM, Brownian motion model; Harvey and Purvis 1991; Butler and King 2004) or if chromosome numbers tend to evolve towards an optimum (OU, Ornstein-Uhlenbeck model; Hansen 1997; Butler and King 2004). We fit BM and OU models on clade-level datasets using the fitContinuous function in the geiger R package (Harmon et al. 2008). We used AIC values (Akaike 1974) for comparing the fit of BM and OU models. Chromosome counts were log-transformed in all phylogenetic comparative analyses.

We also used three non-phylogenetic indices to summarize the patterns of chromosome evolution in animals and plants (Otto and Whitton 2000). First, we calculated the incidence of polyploidy in animals and plants using the distribution of haploid chromosome numbers across species. This measure, also known as the polyploidy index (PI), summarizes the frequency of recent changes across species in haploid chromosome number that have occurred via polyploidization. The PI was estimated as the fraction of all chromosomal changes that involve whole genome duplication as the (#evens - #odds)/#evens. We also determined the significance of the polyploidy index using a binomial test (*sensu* Otto and Whitton 2000). Finally, we estimated the index of support (IS) on the PI to summarize the fraction of the analyzed dataset that must be independent for the PI to remain significantly different from zero. The lower the IS, the stronger the support for a non-zero PI. Second, we estimated the fraction of speciation events associated with changes in chromosome numbers by totaling the minimum number of chromosome changes found within each genus and dividing this by the total number of speciation events within the same genus. Third, we estimated the frequency of speciation events involving polyploidization as the product between the PI and the fraction of speciation events associated with a change in chromosome number. Finally, we compared estimated values for the three indexes between animals and plants (ferns: Otto and Whitton 2000; angiosperms: Rice et al. 2015).

## Results

We present the first database of chromosome counts for the entire animal kingdom. The current version of the ACC includes chromosome counts for 18,349 species across 6,502 genera, 1,320 families, 266 orders, 60 classes, and 21 phyla (Figs. 1, 2). Among the 21 phyla sampled in the current release of the ACC, only seven groups have chromosome counts for ≥1% of their extant diversity: Phoronida (12.5% of total species richness), Chordata (6.5%), Chaetognatha (2.4%), Nematomorpha (2.1%), Sipuncula (1.5%), Entoprocta (1.2%), and Platyhelminthes (1%). Nevertheless, the ACC still lacks of chromosome counts for 11 out of the 27 commonly recognized animal phyla: Acoela, Brachiopoda, Bryozoa, Ctenophora, Gastrotricha, Gnathostomulida, Hemichordata, Kinorhyncha, Onychophora, Placozoa, and Xenoturbellida (Fig. 2). Species in the database have between 1 and 22 haploid counts with an average of 1.14 counts. Chromosome numbers in the current release of the ACC range from *n* = 1–191, with a median chromosome count of 13, a mean of 15.61, and a mode of 12 (1,947 species; Fig. 1).

**Fig. 1.**
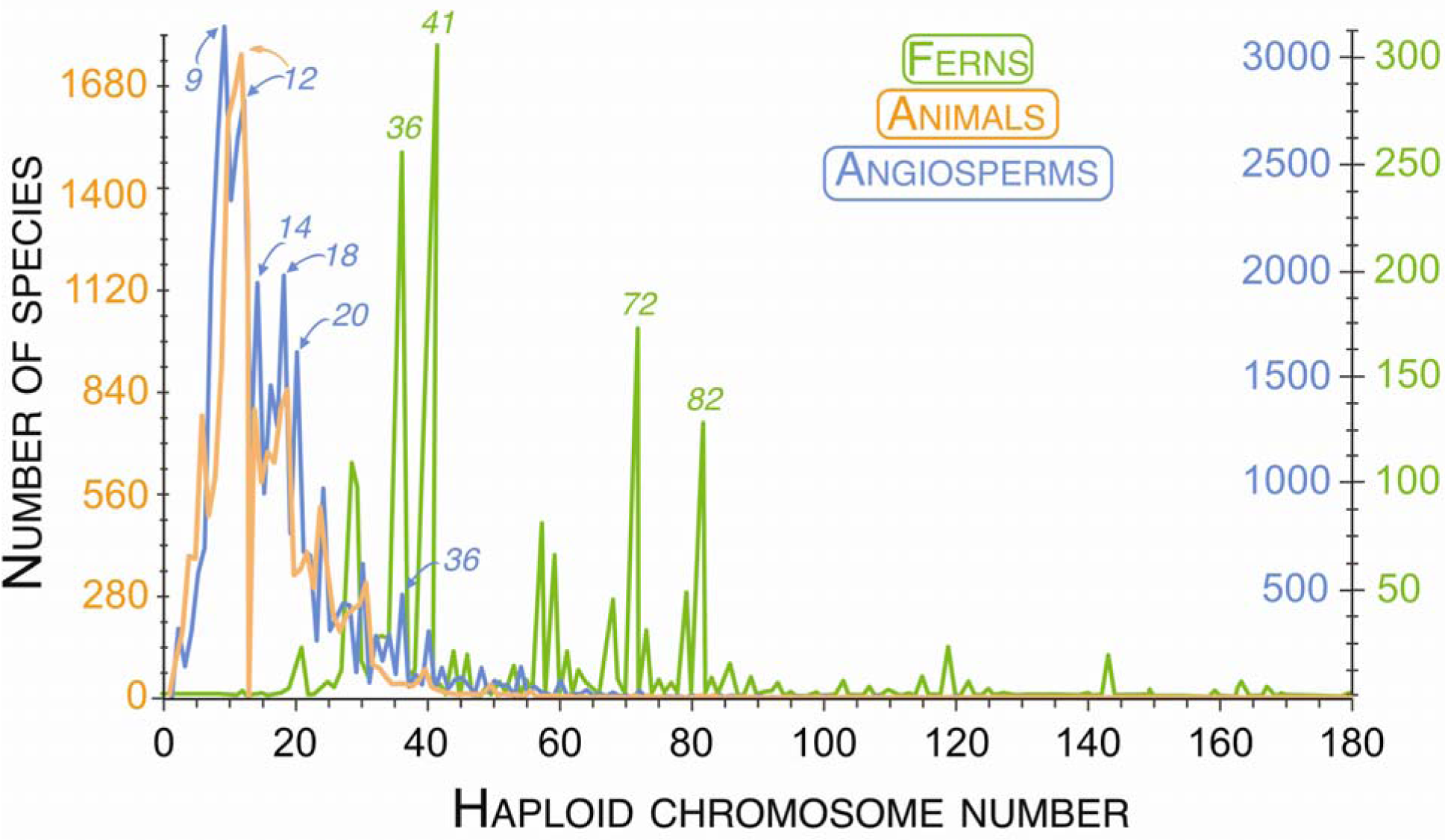
Distribution of haploid chromosome numbers for animals, angiosperms, and ferns. Counts for animals were compiled from the Animal Chromosome Count database (ACC), angiosperms follow the Chromosome Count Database (CCDB, Rice et al. 2015), and the distribution for ferns was modified from Otto and Whitton (2000). Chromosome counts with a relatively high number of species are indicated for each group.

**Fig. 2.**
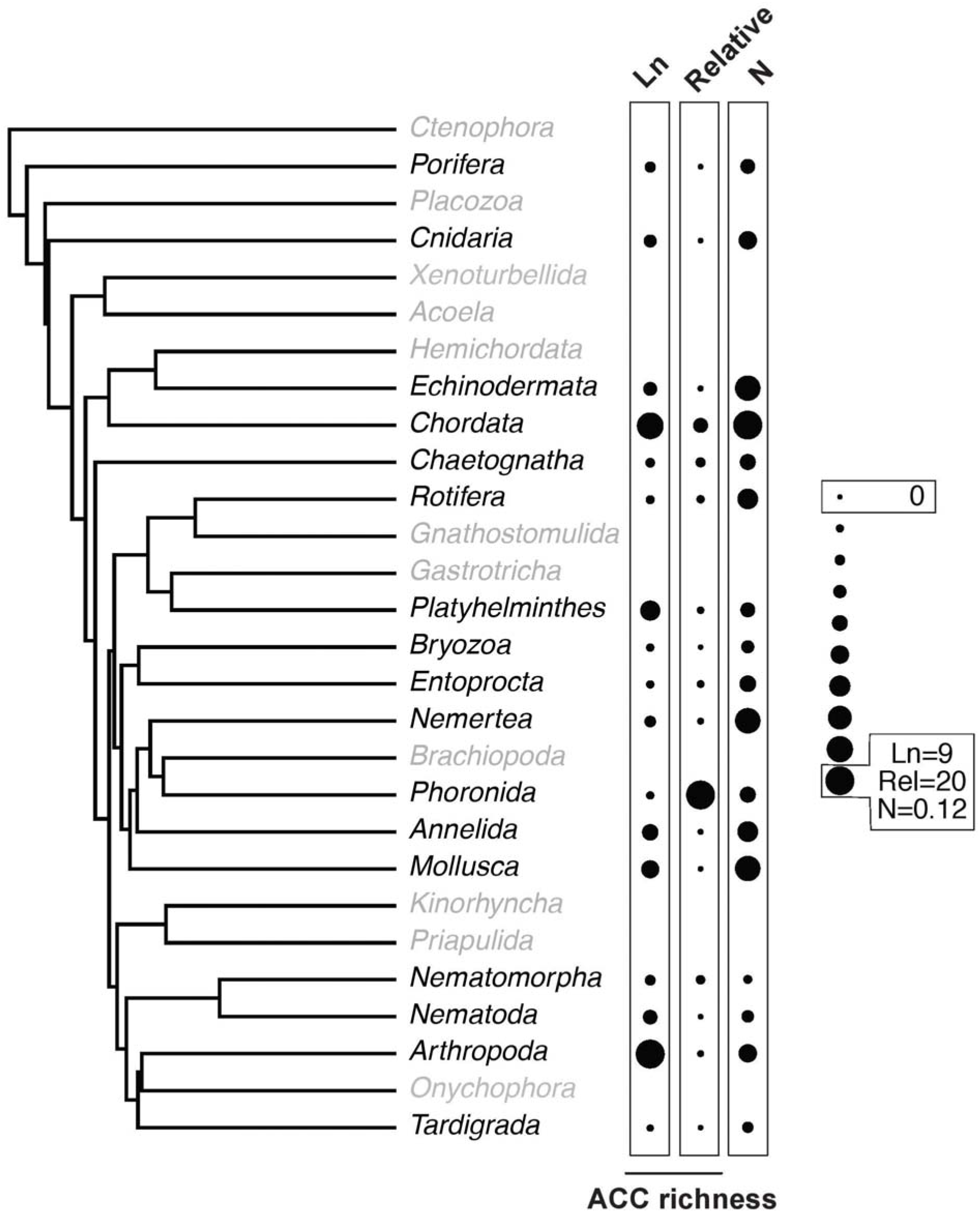
Distribution of the number of species with chromosome counts in the ACC for multiple animal phyla. We show 28 major phyla within animals based on Tree II of Wiens (2015). For each group, we summarize the number of species in the database (Ln column, log-transformed species richness in the ACC), the relative richness in the ACC (number of species divided by the clade richness), and the log-transformed median haploid chromosome number (N). Figure plotted using the phytools R package (Revell 2012). We note that not all animal phyla included in the database are shown in the tree.

Next, we used phylogenetic comparative methods to examine the evolution of chromosome counts across the animal Tree of Life. We found that average differences in median phylum-level chromosome numbers were unrelated to clade age (PGLS *p*-value, Tree I=0.73; Tree II=0.78; Tree III=0.98). Similarly, we found that species richness and chromosome diversity were uncoupled across animal phyla (PGLS Trees I–III, all *p*=0.21). Therefore, across animal phyla, heterogeneity in chromosome numbers does not scale with clade age or richness.

We then tested for phylogenetic conservatism in chromosome numbers within orders of animals and found that patterns of phylogenetic conservatism vary among orders (Fig. 3). Of the analyzed orders, 68% lacked phylogenetic signal in chromosome counts (30 of 44 orders; Fig. 3), including four insect groups (Lepidoptera, Mecoptera, Phasmida, and Psocoptera; Fig. 3), one malacostracan (Isopoda), two actinopterygian fish orders (Siluriformes and Perciformes), two bird clades (Anseriformes and passeriformes), and five mammal clades (Cingulata, Didelphimorphia, Diprotodontia, Perissodactyla, and Soricomorpha).

**Fig. 3.**
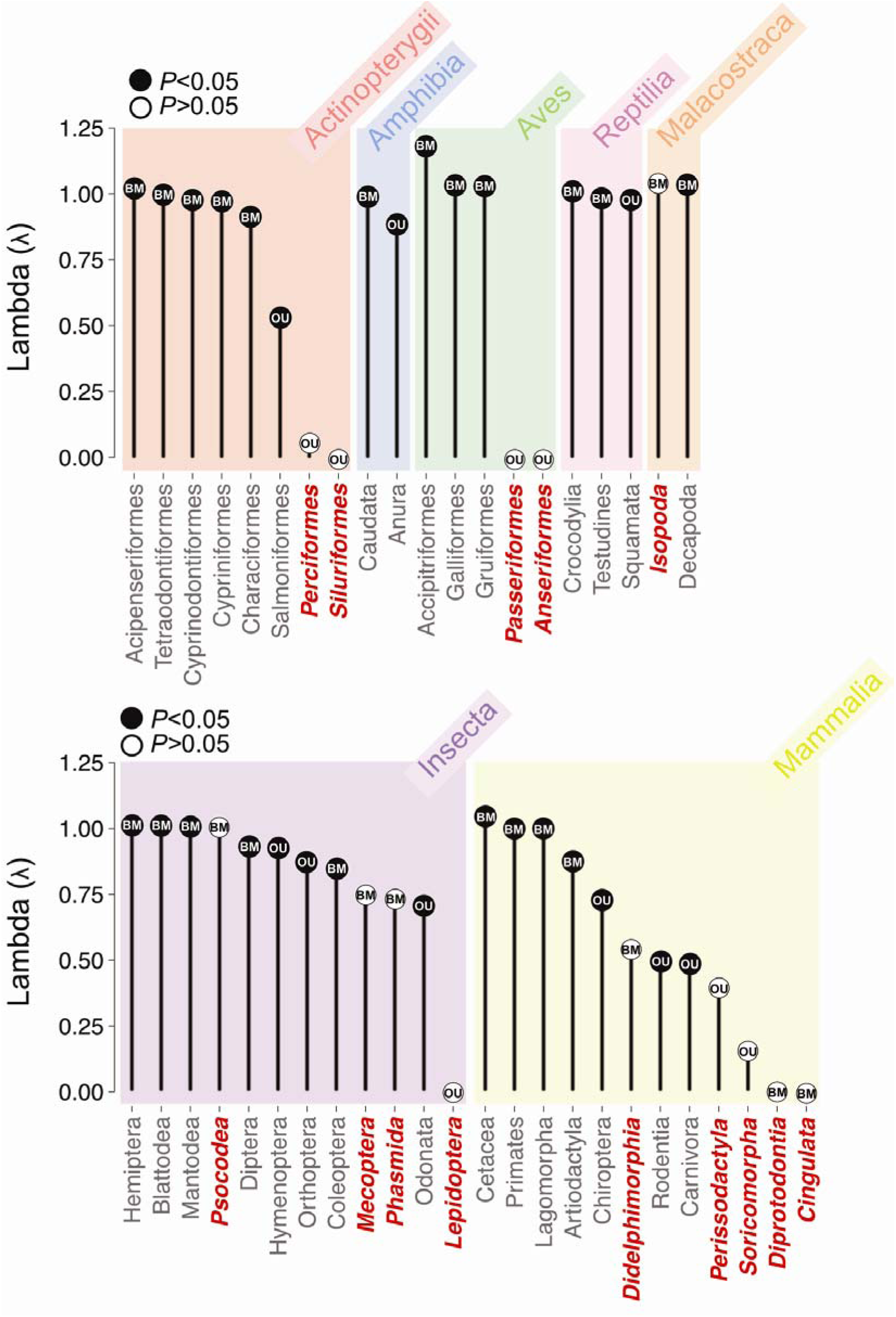
Phylogenetic signal in chromosome counts within animal orders sampled in the ACC. We present estimates of phylogenetic signal in chromosome numbers within each order based on Pagel’s Lambda. We also summarize whether a brownian motion (BM) or Ornstein-Uhlenbeck OU model best fitted the evolution of chromosome numbers across species within each clade. In red, we indicate lineages where no phylogenetic signal was recovered.

Next, we tested whether BM or OU models better explain processes of chromosome number evolution within orders. We found that patterns of chromosome evolution are better explained by OU in 36% of the analyzed orders (16 of 44 orders; Fig. 3) including four insect groups (Hymenoptera, Lepidoptera, Odonata, and Orthoptera), three actinopterygian fish(Perciformes, Salmoniformes, and Siluriformes), a single amphibian clade (Anura), two bird clades (Anseriformes and Passeriformes), five mammal clades (Carnivora, Chiroptera, Perissodactyla, Rodentia, and Soricomorpha), and a single reptilian order (Squamata). Among the clades where OU was the best fit, only 37% (6 of 16) had high and significant phylogenetic signal (lambda > 0.7; P<0.05). Conversely, 75% of the clades where BM was the best fitting model had high and significant phylogenetic signal.

Finally, we compared the evolution of chromosome counts between animals and plants under a non-phylogenetic approach. Different evolutionary processes are responsible for producing similar distributions of chromosome numbers in animals and flowering plants (Fig. 1). For instance, we estimated that 8% of the changes in chromosome number among animals occurred via polyploidization (*PI*=0.0828, *p*<0.001, *IS*=0.6; Table 1) and 34% of the changes in chromosome number occurred with speciation events. In contrast, changes in chromosome number via polyploidization were >3 times more frequent in plants than in animals (angiosperms: *PI*=0.294, *p<*0.001, *IS*=0.1; ferns: *PI*=0.417, *p<*0.001, *IS*=0.039). Similarly, the percentage of speciation events associated with polyploidy was smaller in animals relative to plants (animals: 2.84%; angiosperms: 9.85%; ferns: 7%). However, the frequency of speciation events associated with changes in chromosome number in animals (34%) was similar to the angiosperms (33%) but higher than in ferns (16%).

**Table 1.** Summary statistics for haploid chromosome numbers compiled for ferns, flowering plants, and animals. We summarize the number of species in each database under the # *Species* column. The number of even haploid chromosome counts is indicated under the *# Even* column for each lineage. The percent of species in the database with the modal haploid chromosome count is indicated under the *ModePercent* column. The polyploid index (*PI* column), along with its associated *P*-value, and index of support (*IS* column) are indicated in the table for each lineage. Finally, we summarize for each clade both the fraction of speciation events associated with chromosome changes (*SpeChangeChrom* column) and polyploidization (*SpePolyp* column). We follow Otto and Whitton (2000) for *PI, SpeChangeChrom*, and *SpePolyp* equations and estimates for plants.

| Lineage | # Species | # Even | Mode (%) | PI (%) | P-value [IS%] | SpeChangeChrom (%) | SpePolyp (%) |
| --- | --- | --- | --- | --- | --- | --- | --- |
| Animals | 18343 | 10957 | 0.01 | 8.28 | 1.58E-153 [0.6] | 34.35 | 2.84 |
| Angiosperms | 28199 | 21825 | 11.1 | 29.44 | 5E-324 [0.1] | 33.47 | 9.85 |
| Ferns | 1729 | 1092 | 14.2 | 41.7 | 1E-27 [3.9] | 16 | 7 |

## Discussion

Here we used the most extensive collection of chromosome counts for animals currently available to examine large-scale patterns and processes of chromosome change in animals. Our database contains counts for 18,349 species, representing ∼1% of all described animal species. Although this is the largest and most comprehensive database yet available for animal chromosome counts, many phyla are not represented at all. We still do not know the number of chromosomes—the most basic description of their genomes—for entire clades of animals. These numbers stand in contrast to the information available on plant chromosomes. The plant focused Chromosome Count Database has more than 300,000 chromosome counts for nearly 200,000 species of green plants (Rice et al. 2015). Plant cytology was driven in the early 20th century by the excitement over polyploidy (Barker et al. 2016b), and the result is an extensive catalog of information on plant chromosomes. It is clear from our compilation of data that there is a need for basic cytological research across the animal tree of life. Future releases of the ACC should aim to expand taxonomic sampling within the already sampled phyla, but also, focus on compiling counts for the currently unsampled ones.

### Why is the range of chromosome numbers similar between animals and flowering plants?

We observed a similar distribution in chromosome numbers between animals and flowering plants. Although this could be due to chance, the convergent distribution of numbers suggests that there may be selection for *n*=9–12 chromosomes in both of these lineages. Ferns, in contrast, had much higher numbers of chromosomes with a peak at *n*=41. Chromosome number evolution in the ferns is well recognized as exceptional relative to other eukaryotes (Wood et al. 2009; Otto and Whitton 2000; Barker and Wolf 2010), but the similarity of the distribution of chromosome numbers between flowering plants and animals was not previously recognized. In animals, we found no evidence for an association between chromosome number diversity, clade age, and clade richness across animal phyla that could explain this pattern. Further, variation in chromosome numbers is narrow in many animal subclades, and changes in chromosome number only appear to drive diversity at shorter time scales. Finally, we emphasize that the same haploid number reflects a multitude of different genome organizations among related species. Below, we provide evidence suggesting that there is selection driving a narrow range of chromosome numbers in animals.

First, we found that chromosome numbers do not scale with age and richness at higher taxonomic scales (i.e. across phyla). Species richness and diversity in chromosome numbers are expected to be related if changes in chromosome number are coupled with speciation-related processes (e.g. Lukhtanov et al. 2005; Kandul et al. 2007). Similarly, chromosome diversity should scale with clade age within clades varying in chromosome numbers diversity under a neutral process over time. We found that clade age and species richness do not influence chromosome count heterogeneity across animal phyla. In fact, our results suggest that older clades, which also tend to have more species (e.g. McPeek and Brown 2007), do not exhibit a greater chromosome-level diversity than younger clades. These patterns of time- and richness-uncoupled changes in chromosome number across phyla suggest that chromosome numbers are probably constrained to a certain range at higher taxonomic levels.

Second, we found that changes in chromosome numbers are frequently related to species-level divergence at low taxonomic levels (e.g. within orders). Our results suggest that changes in chromosome numbers may drive patterns of interspecific divergence within some clades.

Specifically, we found an overlap of 66% between clades without phylogenetic signal (*n*=14) and clades where OU was the best fit (*n*=16). Because OU models tend to result in very little phylogenetic signal (Münkemüller et al. 2012), changes in chromosome counts likely played a role in driving species-level divergence within clades such as in Lepidoptera, Perciformes, Siluriformes, Anseriformes, Passeriformes, Perissodactyla, Soricomorpha, Mecoptera, Phasmida, Psocoptera, Isopoda, Cingulata, Didelphimorphia, and Diprotodontia. In addition to agreeing with previous studies on mammals, lepidopterans, and phasmids (Robinson 1971; Bush et al. 1977; White 1978; Scali and Mantovani 1989; Bullini and Nascetti 1990; Passamonti et al. 2004; Lukhtanov et al. 2005; Faria and Navarro 2010; Scali et al. 2012; Potter et al. 2017; Vershinina et al. 2017), our study increases the number of clades where divergence is likely be related to changes in chromosome number.

### Why are changes in chromosome number related to speciation and not polyploidy in animals?

Although animals and flowering plants have similar distributions of chromosome numbers, we found that the underlying processes were different. In particular, polyploidy is more strongly associated with changes in chromosome number and speciation in plants than animals. This is a well known difference in plant and animal speciation (Muller, 1925; Stebbins 1958; Sites and Moritz 1987; Coyne et al. 1993; Otto and Whitton 2000; Gregory and Mable 2005), and our results confirm this long-standing observation. We found that the incidence and importance of polyploidy likely varies between groups within fish, birds, insects, and mammals (Fig. 3). The specific mechanisms explaining why polyploidy is less common in animals probably relate to differences in sex determination (Muller 1925), meiotic disjunction mechanisms (Macgregor 1993), and embryology (von Wettstein 1927; Stebbins 1950), or even the frequency of self-fertilization (White 1973) and the absence of degenerate sex chromosomes (Orr 1990). Ultimately, the relative rarity of polyploidy among animals is still an open question (Gregory 2011), and expanding this database will provide the opportunity for further comparative analyses to test these hypotheses. In contrast, the frequency of speciation-related changes in chromosome number was similar between animals and flowering plants, suggesting that changes in karyotype and chromosome number may have similar impacts on fertility in both lineages.

## Conclusions

The similar distribution of chromosome numbers between animals and flowering plants may be best explained by selection driving numbers toward a narrow range. Our results indicate that the diversity of animal chromosome numbers do not scale with clade age and richness, further suggesting that chromosome numbers are likely under selection across the entire kingdom. Nevertheless, phylogenetic conservatism in chromosome numbers varies across smaller clades in the animal Tree of Life. This variation in constraint on chromosome numbers could be leveraged in future studies to evaluate the causes and consequences of chromosome number evolution. Finally, our analyses indicate that changes in chromosome number are similar between animals and plants. However, polyploidy is more often related to speciation in plants than animals. Although we did not explicitly examine the effects of missing data, future studies based on more extensive species- and population-level datasets should examine, among many others, questions on the association between speciation and chromosomal changes (e.g. Rieseberg 2001), the importance of polyploidy in animals (e.g. Hallinan and Lindberg 2011; Li et al. 2018), and the causes of chromosomal variation among clades and over time (e.g. Martinez et al. 2015; Ross et al. 2015).

## Acknowledgments

Because the ACC is extensively founded on existing chromosome counts compiled by multiple research groups or summarizing counts as a byproduct, we thank the efforts of hundreds of researchers for making data available for others. This database is intended to serve as a quick and highly accessible reference for further research.

## Data accessibility

https://cromanpa94.github.io/ACC/

